# PEDLA: predicting enhancers with a deep learning-based algorithmic framework

**DOI:** 10.1101/036129

**Authors:** Feng Liu, Hao Li, Chao Ren, Xiaochen Bo, Wenjie Shu

## Abstract

Transcriptional enhancers are non-coding segments of DNA that play a central role in the spatiotemporal regulation of gene expression programs. However, systematically and precisely predicting enhancers remain a major challenge. Although existing methods have achieved some success in enhancer prediction, they still suffer from many issues. We developed a deep learning-based algorithmic framework named PEDLA (https://github.com/wenjiegroup/PEDLA), which can directly learn an enhancer predictor from massively heterogeneous data and generalize in ways that are mostly consistent across various cell types/tissues. We first trained PEDLA with 1,114-dimensional heterogeneous features in H1 cells, and we demonstrated that our PEDLA framework integrates diverse heterogeneous features and gives state-of-the-art performance relative to five existing methods for enhancer prediction. We further extended PEDLA to iteratively learn from 22 training cell types/tissues. Our results showed that PEDLA manifested superior performance consistency in both training and independent test sets. On average, PEDLA achieved 95.0% accuracy and a 96.8% geometric mean (GM) across 22 training cell types/tissues, as well as 95.7% accuracy and a 96.8% GM across 20 independent test cell types/tissues. Together, our work illustrates the power of harnessing state-of-the-art deep learning techniques to consistently identify regulatory elements at a genome-wide scale from massively heterogeneous data across diverse cell types/tissues.

## Introduction

Enhancers are distal *cis*-acting DNA regulatory elements that play key roles in controlling cell type-/tissue-specific gene expression ^1–4^ In higher eukaryotes, enhancers recruit transcription factors (TFs) and cofactors to orchestrate vital biological processes including development and differentiation ^5, 6^, maintenance of cell identity ^7–9^, response to stimuli^10–12^, and interactions with target genes through promoter-enhancer looping ^13–16^. Genetic variation or disruption in enhancers is closely associated with diseases and cancers ^17, 18^. Although enhancers have long been recognized for their importance in gene regulation and disease, the absence of common sequence features, their distal location from target genes, and their high cell type/tissue specificity have made them challenging to systematically and precisely identify.

The first genome-scale efforts to identify enhancers were based on evolutionary sequence conservation ^19–22^ because regulatory sequences are likely to evolve under negative evolutionary selection ^23–25^. Owing to the remarkable advances in next-generation sequencing (NGS) technologies, several high-throughput experimental technologies have been developed to predict enhancers in a genome-wide manner. These approaches involve mapping the binding sites of enhancer indicative transcription factors (TFs) ^26–29^ or cofactors ^30–32^ by ChIP-Seq, identifying open chromatin with techniques such as DNase-Seq ^33–35^, and interrogating covalent histone modifications by ChIP-Seq ^7, 12, 36, 37^. Given the diversity and complexity of these high-throughput data, several computational approaches using supervised or unsupervised machine-learning (ML) algorithms have been developed for genome-wide enhancer prediction. Chromia ^38^ uses a hidden Markov model (HMM) to predict regulatory elements, CSI-ANN ^39^ introduces an artificial neural network (ANN) approach, RFECS ^40^ uses random forests to discriminate enhancers from non-enhancers, and DELTA ^41^ uses an adaptive boosting (AdaBoost) approach. ChromaGenSVM ^42^, EnhancerFinder ^43^ and DEEP ^44^ are all based on support vector machine (SVM) classifiers. The unsupervised learning approaches, ChromHMM ^45^ and Segway ^46^, offer genomic segmentation and characterization based on an HMM and a dynamic Bayesian network, respectively.

Although these ML-based computational approaches have achieved remarkable advances in genome-wide identification of enhancers, there remain some critical issues to be settled. First, some of the current computational approaches are limited by a small number of training set samples; thus, they are unlikely to produce a useful general classifier. Second, most of the computational approaches use merely data derived from histone modification marks. Chromia ^38^, CSI-ANN ^39^, RFECS ^40^, DELTA ^41^, ChromaGenSVM ^42^, and DEEP-ENCODE ^44^ combine different numbers and combinations of histone modification marks to predict enhancers. These methods have achieved some success in identifying thousands of enhancers in the human genome. However, it remains a challenging task for these approaches to systematically and precisely predict enhancers, given the simplicity of features derived from only one type of data and the limited number of chromatin modifications that have been examined. Additionally, it is infeasible to fully capture an exhaustive catalogue of chromatin modifications at enhancers. EnhancerFinder ^43^ tackles this issue by simultaneously integrating diverse types of data instead of using a single type of data, thereby improving its developmental enhancer predictions. Third, the class imbalance problem in which the number of enhancer classes is much smaller than the number of non-enhancer classes is extremely common in predicting enhancers ^39, 40, 42, 44^, and ignoring this problem will result in a bias towards the non-enhancer class. RFECS ^40^, DELTA ^41^ and DEEP ^44^ use ensemble techniques to train classifiers with unbalanced classes by combining a set of individual classifiers. Finally, and most importantly, consistent performance across various cell types/tissues is a key challenge in genome-wide enhancer predictions as enhancers are highly cell type-/tissue-specific. The current computational techniques are either trained in a new model for each new cell type/tissue or predict enhancers in new cell types/tissues based on the learned model derived from one cell type/tissue. Regarding the former strategy, this approach is time-consuming and sometimes infeasible given the data availability of training sets for each cell type/tissue, whereas the latter strategy relies on a universal model derived from single cell type/tissue data to effectively identify enhancers in all other cell types/tissues. DEEP-ENCODE ^44^ alleviates this problem by combining four cell-specific models containing 4,000 (4 × 1,000) classifiers in total, and it achieved performance consistency across two other cell lines. However, this is far from sufficient as there exist hundreds and thousands of cell lines and tissue types.

To address these fundamental challenges, we developed a deep learning-based algorithmic framework named PEDLA, which is capable of learning an enhancer predictor that integrates massively heterogeneous data and generalizes in ways that are mostly consistent across various cell types/tissues. We used 1,114-dimensional features in total derived from nine categories of heterogeneous data to predict enhancers in H1 cells. The results show that our PEDLA method has three key advantages: limitless training samples, integration from massively heterogeneous data, and the embedded capability of handling highly class-imbalanced data in an unbiased way. Due to these factors, PEDLA shows state-of-the-art performance relative to five existing methods. Furthermore, we extended PEDLA to iteratively learn from 22 training cell types/tissues from the ENCODE Project ^47^ and the Roadmap Epigenomics Project^48^. The learned optimal model manifested superior classification performance and significant performance consistency in 22 training cell types/tissues and another 20 independent cell types/tissues. Together, our results illustrate the power of harnessing state-of-the-art deep learning techniques to consistently identify regulatory elements on a genome-wide scale from massively heterogeneous data across diverse cell types/tissues.

## Results

### Prediction of enhancers using PEDLA with heterogeneous signatures

To systematically and precisely predict enhancers on a genome-wide scale, we developed a deep learning-based algorithm framework named PEDLA (see Materials and Methods, Fig. S1). Briefly, we first used nine categories of heterogeneous data in H1 cells that serve as discriminating features to identify enhancers, including histone modifications (ChIP-Seq), TFs and cofactors (ChIP-Seq), chromatin accessibility (DNase-Seq), transcription (RNA-Seq), DNA methylation (RRBS), CpG islands, evolutionary conservation, sequence signatures, and occupancy of TFBSs. In total, 1,114-dimensional features were used in the training and testing of PEDLA (Table S1). Then, we constructed the enhancer class (positive class set) containing 5,870 enhancer regions (Table S2) based on H3K27ac peaks in the H1 cell line as no 'gold standard' set of enhancers has been experimentally verified across various cell types/tissues. Next, we maintained a ratio of 1:10 between positive and negative samples by choosing an equal number of promoters and nine times the number of random background regions that were not annotated as promoters or enhancers as the non-enhancer class (negative class set). Finally, we optimized the structure of PEDLA by selecting the most optimal model, which consists of two hidden layers with each layer including 500 hidden units, in terms of accuracy using 5-fold cross-validation in both the training set and test set (Fig. 1A, Table S3). On average, PEDLA achieved 97.7% accuracy, a 97.2% GM (96.6% sensitivity and 97.9% specificity) and an 88.7% F1-score (96.6% recall and 82.1% precision) for the training set, as well as 97.7% accuracy, a 97.0% GM (96.2% sensitivity and 97.8% specificity) and an 88.3% F1-score (96.2% recall and 81.7% precision) for the independent test set. These results demonstrated the superior performance and outstanding robustness of PEDLA in both the training and test sets, suggesting that PEDLA has great representational power for heterogeneous data and excellent ability to solve the overfitting problem.

**Figure 1.**
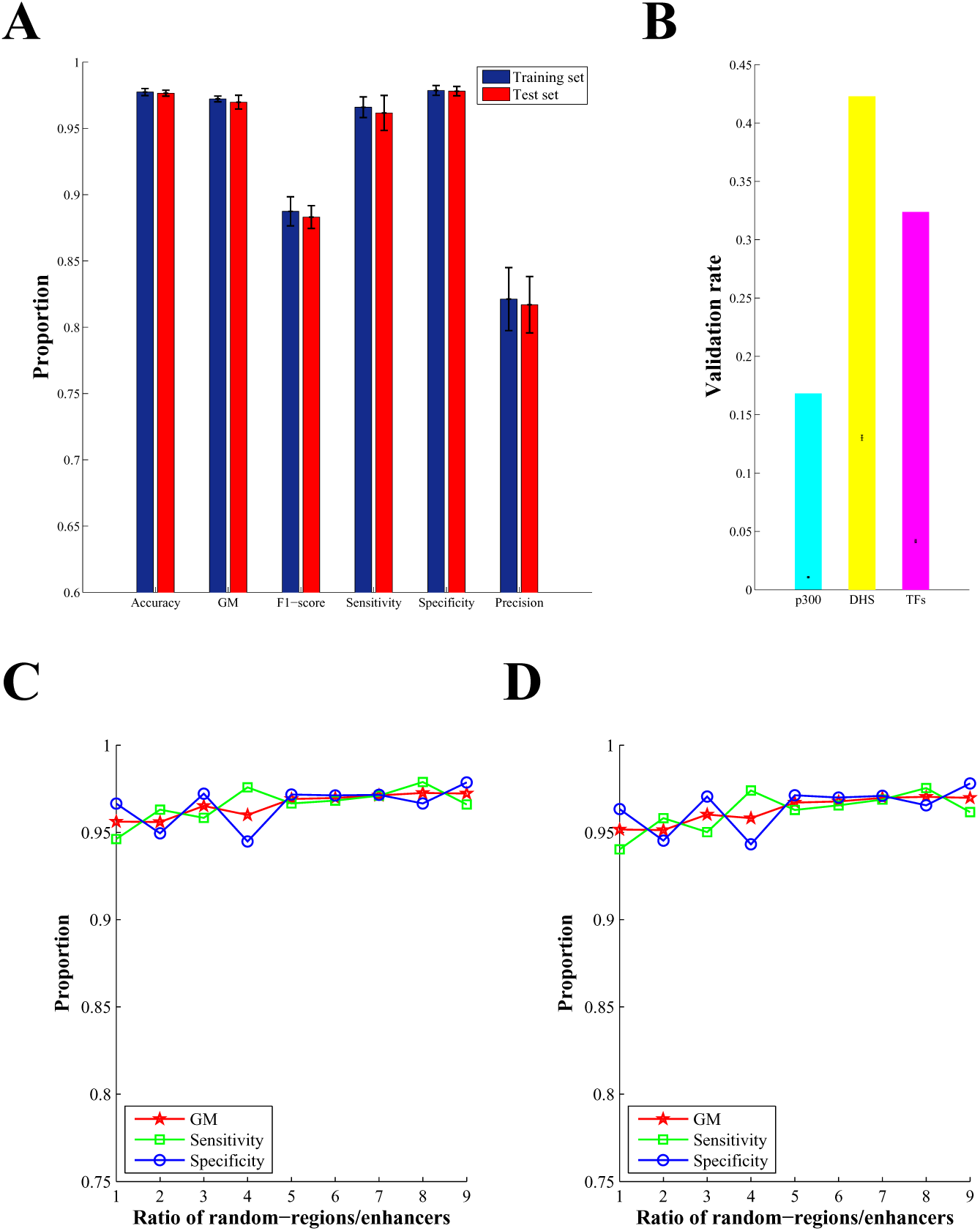
Enhancer predictions using PEDLA with heterogeneous signatures and class-imbalanced data in H1 cells. (A) Selecting the optimal structure of PEDLA for the purpose of enhancer prediction using 1,114 heterogeneous signatures. Three performance indicators, accuracy, GM and F1-score, were measured using 5-fold cross-validation in both a training set and test set. The error bar indicates the mean and standard deviation of the performance indicator. (B) Validation of enhancer predictions by distal DHSs, binding sites of p300 and TFs (NANOG, OCT4 and SOX2) using the trained PEDLA with optimal structure. The bar shows the actual validation rate, whereas the error bar shows the mean and standard deviation of validation rates for 10,000 randomly shuffled predictions. (C-D) Capability of handling class-imbalanced data unbiasedly. Three performance indicators, sensitivity, specificity and GM, were measured for the training set (C) and test set (D) using 5-fold cross-validation based on the optimal structure of PEDLA with all 1,114-dimensional features. The number of enhancers, promoters and random regions not annotated as promoters or enhancers were maintained at 1 : 1 : *x* (*x* = 1, 2, …, 9), such that the ratio between positive and negative samples was 1 : (1 + *x*).

### Ability to handle class-imbalanced data in an unbiased manner

To resolve the class imbalance problem, our PEDLA method has an embedded mechanism in which the prior probability ***P*** of each class, which reflects the imbalance of the class data, is directly estimated from the training data. Dividing the posterior probability by the prior probability *P* can fundamentally eliminate class-imbalanced influence (see Materials and Methods, Fig. S1). Thus, for PEDLA, we need to train only a single model instead of combining multiple trained classifiers as ensemble approaches do, which generally results in a bloated structure model. To assess the performance of PEDLA in resolving this issue, we maintained the number of enhancers, promoters and random regions not annotated as promoters or enhancers in the training set of the H1 cell line at 1 : 1 : *x* (*x* = 1, 2, …, 9), such that the ratio between positive and negative samples was 1 : (1 + *x*). Three performance indicators, namely sensitivity, specificity and GM, which were reported to be suitable for assessing highly imbalanced data sets, were measured using 5-fold cross-validation based on the optimal model of PEDLA with all 1,114-dimensional features (Fig. 1C–D). For comparison, we additionally developed a classic DNN algorithm and assessed the performance of the classic DNN in resolving this challenge by measuring the three performance indicators (Fig. S2). We found that for PEDLA, all three performance indicators were unbiased for different degrees of imbalance of the class-imbalanced data in both the training and test data. Notably, the performance of PEDLA even showed a certain degree of improvement with increasing imbalance. In contrast, for the classic DNN, the sensitivity and GM performance indicators decreased rapidly as the specificity increased slightly with increasing imbalance of the class data. The higher specificity and the lower sensitivity achieved by the classic DNN for highly class-imbalanced data suggested that the classic DNN tended to be biased towards the majority class, which is similar to the individual SVM for imbalanced data ^49^. These results indicate that PEDLA manifested greater capability in resolving the class imbalance problem.

### Validation of predicted enhancers

To validate the predicted enhancers using PEDLA, we calculated the validation rate as the percentage of predicted enhancers overlapping with experimental data of enhancer markers, including distal DHSs, p300 binding sites and a few sequence-specific TFs known to function in H1 cells, such as NANOG, OCT4 and SOX2 (see Materials and Methods). Additionally, we computed the misclassification rate as the percentage of predicted enhancers overlapping with annotated promoters (see Materials and Methods). In human H1 cells, we identified 20,689 enhancers using the trained PEDLA with optimal structure, and we found that 42.3%, 16.8%, and 32.4% of these predicted enhancers overlapped with distal DHSs and binding sites of p300 and TFs, respectively. With 10,000 iterations of random shuffling of the enhancer predictions in the H1 genome, we found the average validation rate of distal DHS, p300 and TF to be 13.0%, 1.1% and 4.2%, respectively, and the actual validation rates were highly significant according to a one-sided *t*-test (*p*-values = 0) (Fig. 1B). Additionally, we found that 6.6% of enhancer predictions overlapped with TSSs annotated by GENCODE (V15) ^50^, indicating a low misclassification rate. These results demonstrate that PEDLA can accurately predict putative enhancers based on the nine categories of heterogeneous data.

### Performance comparisons of PEDLA with existing methods

Next, we compared our enhancer predictions with those identified by five state-of-the-art methods, including the supervised learning methods CSI-ANN ^39^, RFECS ^40^, and DELTA ^41^ and the unsupervised learning methods ChromHMM ^45^ and Segway ^46^, in H1 cells. The optimal model of CSI-ANN was obtained based on 39 histone modifications from CD4+ T cells ^39^, the best model of RFECS was derived from the profiles of 24 histone modifications in H1 and IMR90 cells ^40^, whereas the optimal model of DELTA was trained on the optimal subset of histone modifications derived from H1 and CD4+ T cells ^41^. To make a fair comparison of performance of our method with those of the three supervised approaches, we applied PEDLA and the three supervised learning methods to a common histone modification dataset in H1 cells. For the common dataset, we used H3K4me1, H3K4me3 and H3K27ac as the minimal chromatin marks for enhancer prediction as these three histone markers were previously used as the minimal subset of histone modifications to predict enhancers by both CSI-ANN ^39^ and RFECS ^40^ and were also ranked within the top 4 in terms of variable importance in DELTA in H1 cells ^41^. Using these three histone markers in H1 cells, RFECS, CSI-ANN, and DELTA made 75,084, 30,173, and 112,044 enhancer predictions, respectively, based on the optimal parameters and default threshold used in their studies. PEDLA yielded 22,691 enhancers using the optimal structure of a hidden layer with 50 hidden units determined by 5-fold cross-validation. For ChromHMM and Segway, the enhancer predictions can be directly downloaded and extracted from the ENCODE page (http://hgdownload.cse.ucsc.edu/goldenPath/hg19/encodeDCC/wgEncodeAwgSegmentation/), and obtained 26,869 and 131,698 predictions, respectively.

We compared PEDLA with the five existing methods based on nine performance indicators: accuracy; sensitivity; specificity; GM; F1-score; validation rates of distal DHSs, P300, and TFs; and misclassification rate. The comparative analysis across these methods is summarized in Table 1. Based on the performance indicators accuracy, sensitivity, GM, F1-score, and validation rate (DHS, P300, and TFs), PEDLA consistently performed better than all other methods. Based on specificity, two unsupervised methods (ChromHMM and Segway) were ranked first followed by CSI-ANN, RFECS, PEDLA and DELTA. Based on misclassification rate, DELTA and RFECS shared the best results, followed by ChromHMM, PEDLA, Segway and CSI-ANN. Furthermore, comparing PEDLA of the optimal model based on the three minimal chromatin marks with that based on the full 1,114 features, we found that PEDLA of full features consistently performed better than PEDLA of the three minimal chromatin marks for all performance indicators. This result suggests that approaches integrating diverse types of data give more complete and precise representation of enhancers and thus can significantly improve enhancer prediction compared to methods using only a single type of data.

**Table 1.**
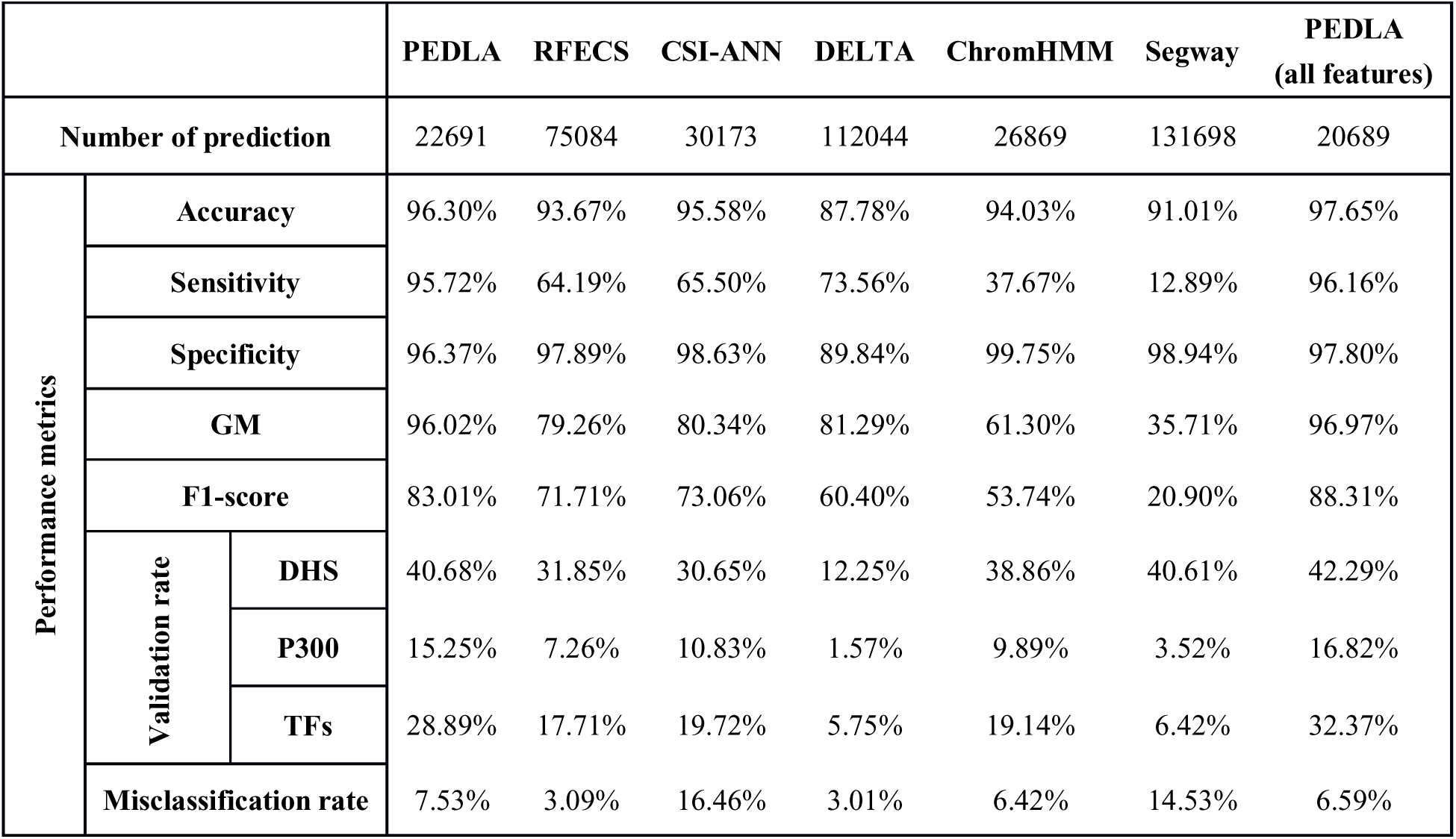
Comparison of the performance of PEDLA with that of existing methods

Because the different performance indicators illustrate the distinct advantages and disadvantages of these studied methods, we ranked their performance according to the nine metrics. In total, we performed nine different tests, including seven methods and nine performance indicators. According to a previously published method ^51^, we averaged the ranked positions of each of the seven methods used in comparison in all nine tests. Table 2 demonstrates the overall score and average rank position of each of the methods; a lower average rank indicates better performance. This analysis revealed that across the different performance tests, the PEDLA with both full signature sets and three minimal chromatin marks shared the best results, followed by CSI-ANN, ChromHMM, RFECS, DELTA and Segway.

**Table 2.** Relative ranking of PEDLA and existing methods based on the results of our comparison study

| **Performance metrics** |  | **PEDLA** | **RFECS** | **CSI-ANN** | **DELTA** | **ChromHMM** | **Segway** | **PEDLA**<br>**(all features)** |
| --- | --- | --- | --- | --- | --- | --- | --- | --- |
| **Accuracy** |  | 2 | 5 | 3 | 7 | 4 | 6 | 1 |
| **Sensitivity** |  | 2 | 5 | 4 | 3 | 6 | 7 | 1 |
| **Specificity** |  | 6 | 4 | 3 | 7 | 1 | 2 | 5 |
| **GM** |  | 2 | 5 | 4 | 3 | 6 | 7 | 1 |
| **F1-score** |  | 2 | 4 | 3 | 5 | 6 | 7 | 1 |
| **Validation rate** | **DHS** | 2 | 5 | 6 | 7 | 4 | 3 | 1 |
|  | **P300** | 2 | 5 | 3 | 7 | 4 | 6 | 1 |
|  | **TFs** | 2 | 5 | 3 | 7 | 4 | 6 | 1 |
| **Misclassification rate** |  | 5 | 2 | 7 | 1 | 3 | 6 | 4 |
| **OVERALL RANKING** |  | 2nd (25), 2.78 | 5th (40), 4.44 | 3rd (36), 4.00 | 6th (47) 5.22 | 4th (38), 4.22 | 7th (50), 5.56 | 1st (16), 1.78 |

### Further performance assessment across methods

To further assess the performance across these supervised methods, we should exclude the bias of different numbers of enhancer predictions of these methods on performance comparison. Thus, we selected thresholds that yielded comparable numbers of predictions for RFECS, CSI-ANN, and DELTA, so as to make a fair comparison across these methods. We obtained 20,595, 19,859, and 20,679 enhancer predictions for RFECS, CSI-ANN, and DELTA, respectively, which were comparable to the number of predictions obtained by our PEDLA, 22,691 (Table 3). Comparing with the data presented in Table 1, we found that the three methods achieved higher validation rate and lower misclassification rate, but they obtained a much lower accuracy. Most importantly, the sensitivity, GM and F1-score of these three methods decreased rapidly. These results suggested that the performance comparison analysis at comparable numbers of predictions across these methods seems fair, however, such analysis might deflate the performance of the three methods in some cases. Solely comparing performance at similar numbers of enhancer predictions across methods might sacrifice the balance between different performance indicators.

**Table 3.** Performance comparisons across methods with considerable enhancer predictions

|  |  |  | **PEDLA** | **RF ECS** | **CSI-ANN** | **DELTA** | **PEDLA**<br>**(all features)** |
| --- | --- | --- | --- | --- | --- | --- | --- |
| **Number of prediction** |  |  | 22691 | 20595 | 19859 | 20679 | 20689 |
| **Performance metrics** | **Accuracy** |  | 96.30% | 91.83% | 92.42% | 88.80% | 97.65% |
|  | **Sensitivity** |  | 95.72% | 36.92% | 20.07% | 12.81% | 96.16% |
|  | **Specificity** |  | 96.37% | 99.68% | 99.74% | 99.84% | 97.80% |
|  | **GM** |  | 96.02% | 60.65% | 43.73% | 35.37% | 96.97% |
|  | **F1-score** |  | 83.01% | 53.03% | 31.92% | 22.28% | 88.31% |
|  | **Validation rate** | **DHS** | 40.68% | 38.40% | 28.66% | 11.25% | 42.29% |
|  |  | **P300** | 15.25% | 15.03% | 12.34% | 2.93% | 16.82% |
|  |  | **TF** | 28.89% | 25.50% | 18.84% | 8.13% | 32.37% |
|  | **Misclassification rate** |  | 7.53% | 1.73% | 20.96% | 1.43% | 6.59% |

All the trainings of these supervised methods were based on a positive enhancer set (5,870 enhancer regions, Table S2) that were constructed by calling H3K27ac peaks in the H1 cell line using MACS ^52^ with a stringent significance level (*p*-value < 10^−9^). To test whether the positive enhancer set has effects on the performance assessment of these methods, we constructed two independent positive enhancer sets by calling H3K27ac peaks using two less stringent significance levels (*p*-value < 10^−6^ and *p*-value < 10^−4^). We obtained 8,413 and 11,937 enhancers as positive set, and re-trained the models of these methods with these two less stringent training sets, respectively. We summarized the results of performance assessment across methods in Table 4 and Table S6, respectively. Our results suggested that in both cases, our PEDLA achieves much better performance relative to other methods. To further strengthen our finding, we re-trained the models of these methods with the positive enhancer set provided by the study of RFECS ^40^, which consists of 5,899 active and distal p300 binding sites (Table 5). Our results demonstrated that our PEDLA also illustrates superior performance comparing with other methods. Taken together, our results convincingly demonstrate that PEDLA integrates diverse heterogeneous features and gives superior performance relative to the existing methods for enhancer predictions.

**Table 4.** Performance comparisons across methods with less stringent positive enhancer sets *p*-value < 10^−6^

|  |  |  | **PEDLA** | **RF ECS** | **CSI-ANN** | **DELTA** | **PEDLA**<br>**(all features)** |
| --- | --- | --- | --- | --- | --- | --- | --- |
| **Number of prediction** |  |  | 32035 | 31779 | 31827 | 31837 | 31238 |
| **an** | **ce** | **Accuracy** | 97.07% | 92.78% | 93.24% | 89.72% | 96.59% |
|  | **Sensitivity** |  | 93.41% | 46.71% | 30.03% | 21.00% | 96.92% |
|  | **Specificity** |  | 97.45% | 99.36% | 99.63% | 99.71% | 96.56% |
|  | **GM** |  | 95.39% | 68.11% | 54.49% | 45.57% | 96.74% |
|  | **F1-score** |  | 85.53% | 61.78% | 44.67% | 33.97% | 84.01% |
|  | **Validation**<br>**rate** | **DHS** | 38.94% | 35.18% | 30.81% | 11.03% | 39.68% |
|  |  | **P300** | 12.29% | 11.73% | 10.59% | 2.81% | 13.07% |
|  |  | **TF** | 25.97% | 21.73% | 19.68% | 7.82% | 29.14% |
|  | **Misclassification rate** |  | 6.81% | 2.00% | 15.89% | 1.48% | 6.71% |

**Table 5.**
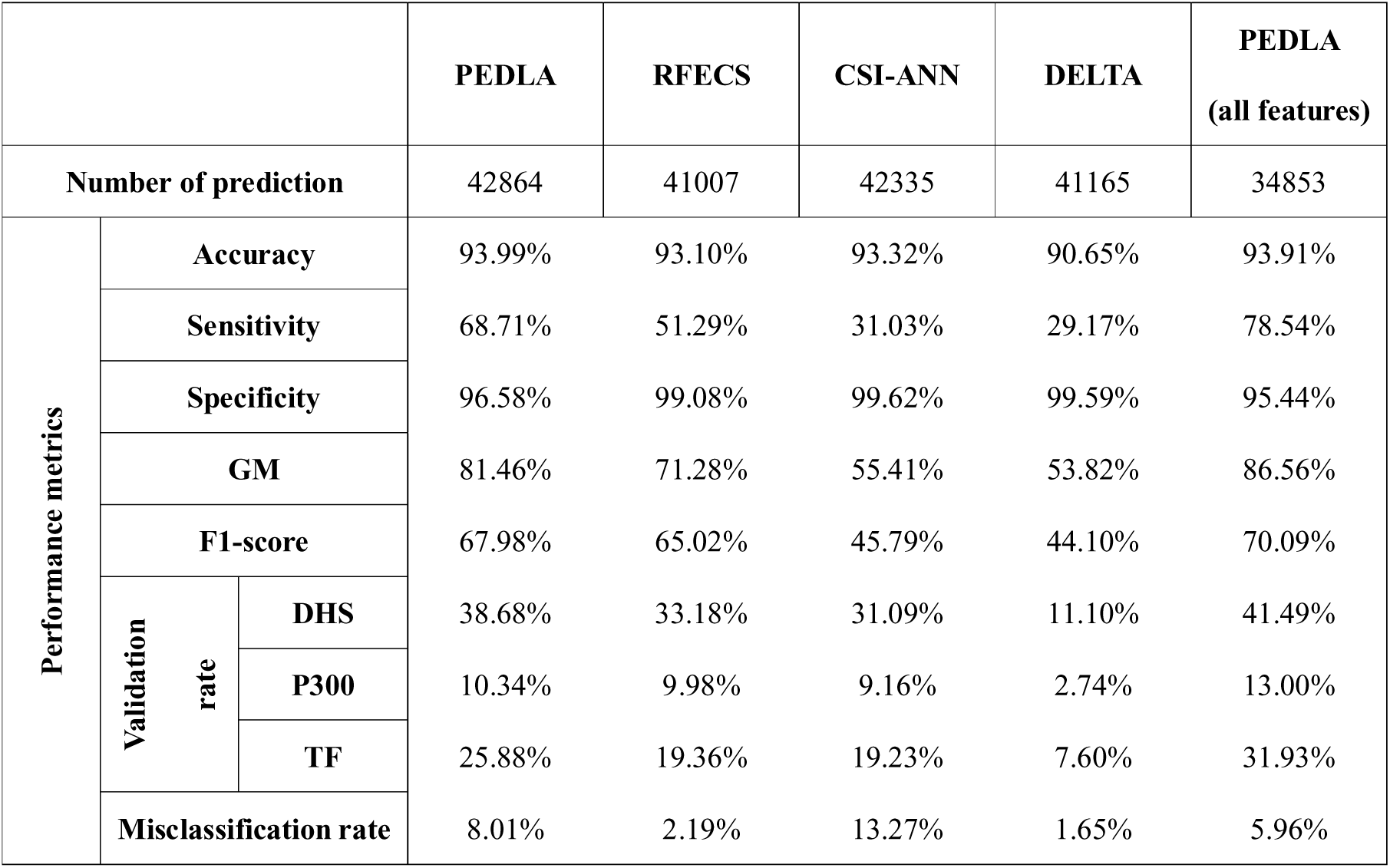
Performance comparisons across methods with positive enhancer sets provided by RFECS

### Prediction of enhancers with PEDLA in multiple human cell types

Predicting enhancers across more human cell types/tissues can help us better understand the underlying regulatory mechanisms of these elements. Currently, most computational approaches use individual models trained on data from one cell type/tissue to predict enhancers in multiple other cell types/tissues. Because enhancers are highly cell type-/tissue-specific, it is infeasible that a model derived from single cell type/tissue data can effectively predict enhancers in all the other types/tissues. To resolve this challenge, DEEP-ENCODE ^44^ combines four ensemble models that are trained on four cell lines and contain 4,000 (4 × 1000) classifiers in total to predict enhancers in multiple cell types, and the test results in two independent cell lines suggested that DEEP-ENCODE showed better generalization capabilities. However, to achieve consistent performance across various cell types/tissues, DEEP-ENCODE has to combine many cell type-/tissue-specific models, which makes the structure of DEEP-ENCODE more bloated.

Inspired by the core idea of deep learning using unsupervised pre-training followed by supervised fine-tuning, we developed an iterative PEDLA for enhancer prediction across diverse human cells and tissues to resolve the above-mentioned issues. Figure 2 shows a diagram of the steps of training PEDLA in multiple cell types/tissues. Initially, PEDLA was trained based on data derived from any cell type/tissue. Then, PEDLA was trained for a subsequent cell type/tissue iteratively, using the trained model of the previous cell type/tissue as initialization. For each training iteration, PEDLA learned cell type-/tissue-specific enhancer information from the corresponding cell type/tissue and “evolved” iteratively. Finally, PEDLA learned enhancer information derived from all the cell types/tissues used in the training, and it showed consistent performance in predicting enhancers across diverse cell types/tissues in the human genome.

**Figure 2.**
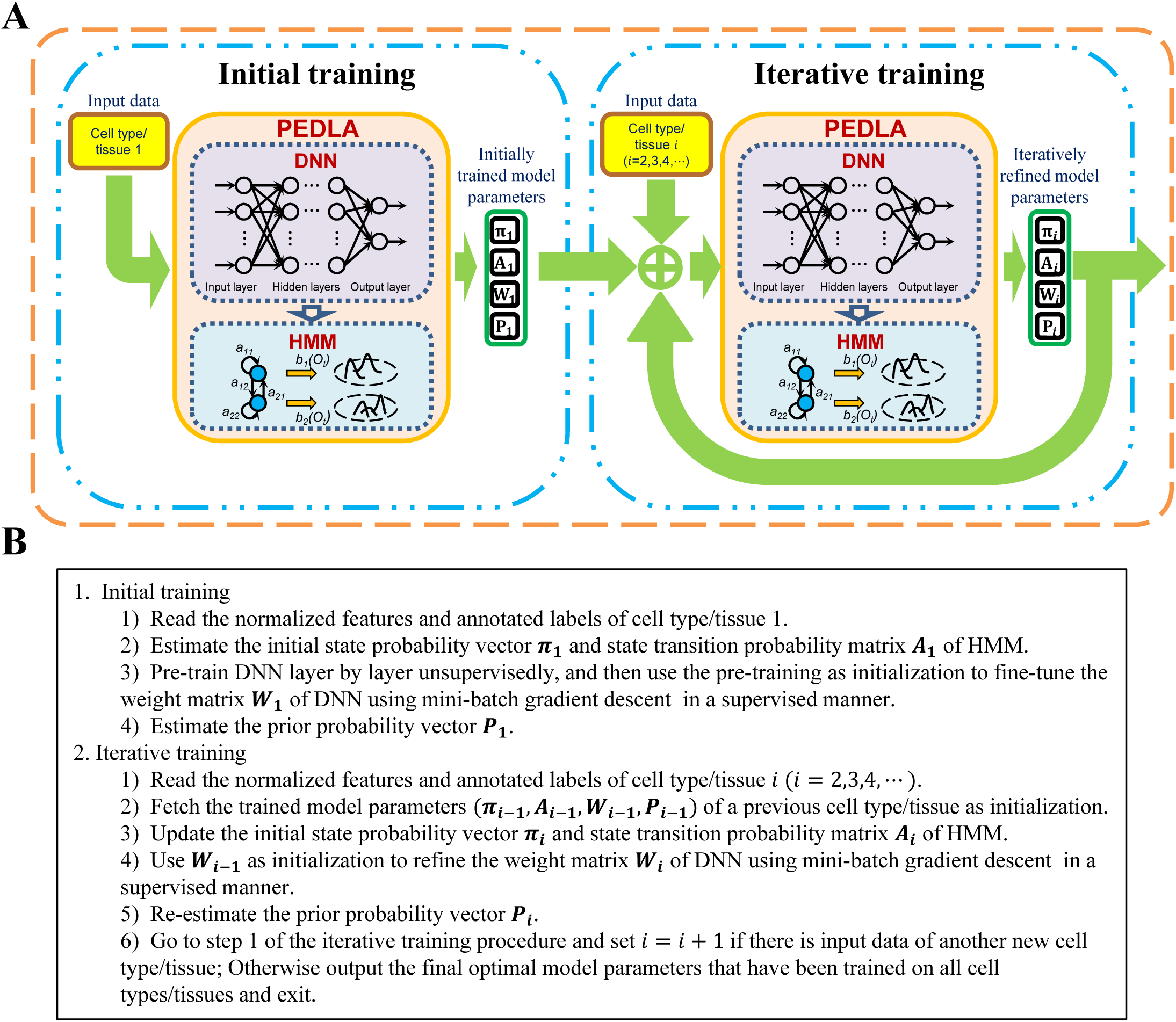
Training procedure of PEDLA in multiple human cell types. (A) Schematic diagram showing the framework of training PEDLA for the purpose of enhancer prediction in multiple human cells/tissues. The whole training procedure comprises initial training and iterative training. (B) The pseudocode shows the detailed steps of training PEDLA in multiple human cells/tissues.

To explore the effectiveness of PEDLA to predict enhancers in multiple cell types/tissues, we selected 22 cell types/tissues from the ENCODE Project ^47^ and the Roadmap Epigenomics Project ^48^ as the training cell set, and we chose another 20 independent cell types/tissues that were not used in the training procedure as the test cell set (Table S4). The training and test procedures of PEDLA are much more time-consuming when more features are used. Thus, we used the 8-dimensional features, including H3K4me1, H3K4me3, H3K9me3, H3K27me3, H3K36me3, H3K27ac, input of histone modification and evolutionary conservation as the common features to train PEDLA across multiple cell types/tissues. The selection of these features is based on the consideration of both the data availability across tens of cell types/tissues and the variable importance of all features in enhancer predictions (Table S5), which was assessed using R package called randomForest ^53^ based on average accuracy.

PEDLA was trained based on the enhancer and non-enhancer classes that were constructed independently for each of the 22 cell types/tissues in the training cell sets using the optimal model of a hidden layer with 50 hidden units (Fig. 3A). The whole training procedure was repeated with 50 random permutations of the 22 training cell types/tissues (50 out of *P*(22, 22) = 22!), and each random order was repeated four times with random permutations of training samples for each cell type or tissue. In total, the training of PEDLA was repeated independently 200 times on the 22 training cell types/tissues. During each iterative step of the training procedures, we used the optimal model of PEDLA to predict enhancers in the 22 training cell types/tissues and to simultaneously assess performance on another 20 independent test cell types/tissues (Fig. 3B–C). We recorded the three performance indicators (accuracy, GM, and F1-score) across these cell types/tissues, which made it intuitive to interrogate the dynamic performance of PEDLA with the increasing number of cell types/tissues that it had been trained on. We found that the mean values of the performance indicators increased and that the variances of the performance indicators decreased with an increasing number of training cell types/tissues, suggesting that increased performance and increased consistency were obtained for of our PEDLA enhancer predictions, respectively. Furthermore, the three performance indicators increased rapidly when the second cell type/tissue was added into the training, indicating that the performance and consistency of PEDLA were greatly improved compared to the performance and consistency of the PEDLA that was trained on only one cell type/tissue. After finishing the training of the five cells/tissues, the performance indicators remained unchanged, suggesting that the performance and the consistency of PEDLA were stable at this stage.

**Figure 3.**
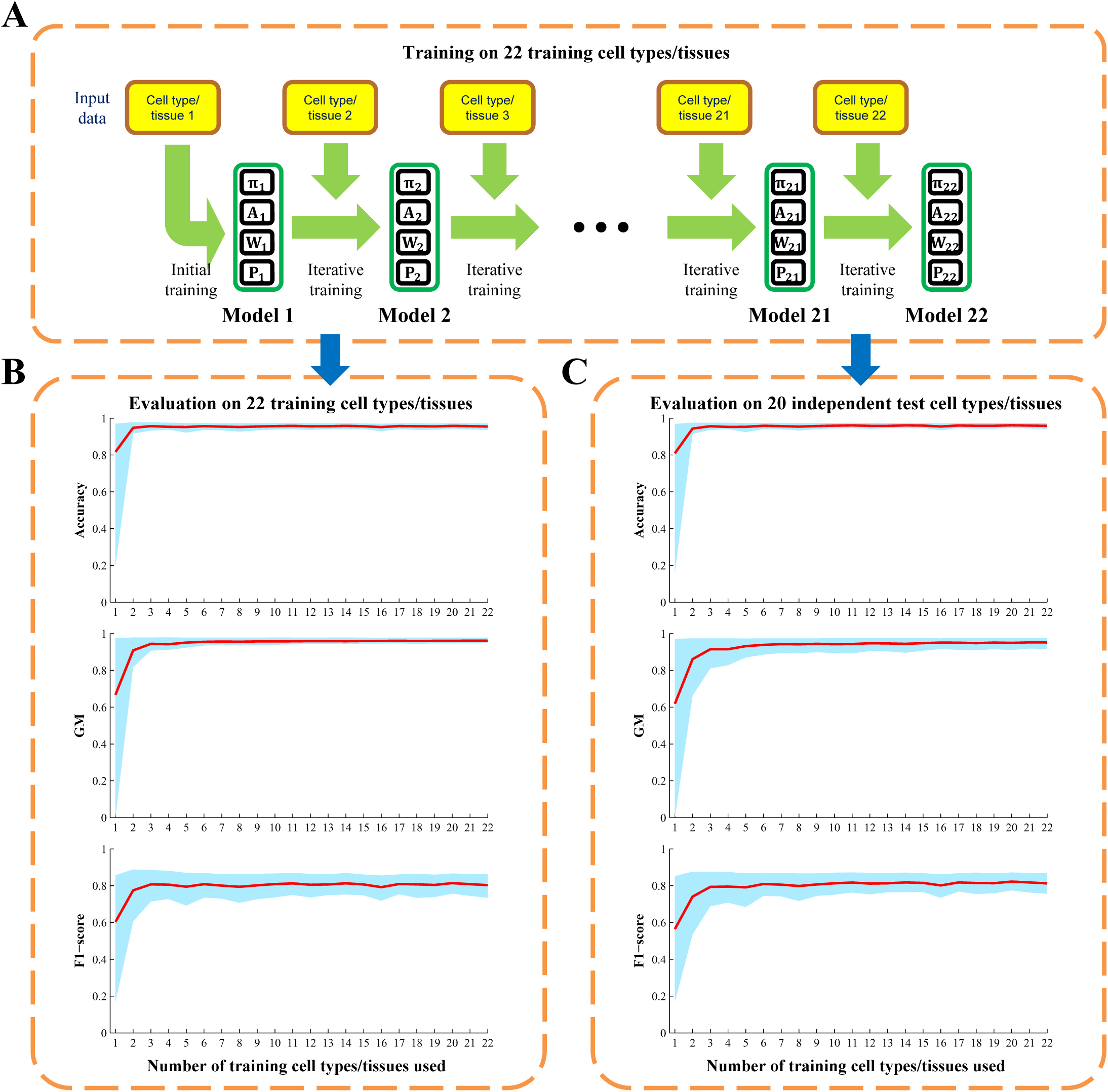
Evaluation of enhancer predictions using PEDLA in multiple human cells/tissues. (A) Evaluation of enhancer prediction using PEDLA in 22 training cell types/tissues. The whole training procedure was repeated with 50 random orders of the 22 training cell types/tissues, and each random order was repeated four times with random permutations of training samples for each cell type/tissue. In total, the training of PEDLA was repeated 200 times on the 22 training cell types/tissues, independently. For each repeat, the trained optimally model of PEDLA was saved for later evaluation for each of the 22 training cell types/tissues. Thus, 22 × 200 = 4,400 optimal models were generated. 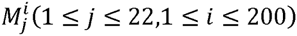 denotes the optimal model that finished training on the *j*-th training cell type/tissue in the *i*-th run of the 200 independent runs. (B-C) Performance evaluations of PEDLA in the training cell set and the independent test cell set along the training route. Three performance indicators, accuracy, GM, and F1-score, were assessed for the PEDLA with the trained optimal model in the 22 training cell types/tissues and 20 test cell types/tissues, independently. (B) For a fixed *j* of the X-axis, all 200 optimal models 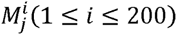 were used to assess the performance indicators on the 22 training cell types/tissues. The red line represents the mean of the total 200 × 22 = 4,400 values of each performance indicator, and the light blue colour band indicates the 10^th^ and 90^th^ percentiles. (C) For a fixed *j* of the X-axis, all 200 optimal models 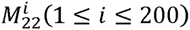 were used to assess the performance indicators on the 20 test cell types/tissues. The red line represents the mean of the total 200 × 20 = 4,000 values of each performance indicator, and the light blue colour band indicates the 10^th^ and 90^th^ percentiles.

We use the best-trained model of PEDLA that had been completed using the training on all 22 training cell types/tissues to further assess its generalizability using the three performance indicators (Fig. 4; Table S7-8). On average, PEDLA achieved 95.0% accuracy, a 96.8% GM (99.0% sensitivity and 94.6% specificity) and a 78.7% F1-score (99.0% recall and 65.4% precision) for 22 training cell types/tissues, as well as 95.7% accuracy, a 96.8% GM (98.2% sensitivity and 95.4% specificity) and an 81.0% F1-score (98.2% recall and 69.1% precision) for 20 independent test cell types/tissues. This result suggests that PEDLA shows superior performance and consistency across various cell types/tissues and generalizes well to both the training and test type/tissue sets. To further verify the performance consistency of PEDLA across diverse cell types/tissues, we assessed the performance of PEDLA for predicting specific and common enhancers across the cell types/tissues in both the training set and the test set (Fig. 4; Table S7-8). In the 22 training cell types/tissues sets, PEDLA achieved 94.9% accuracy, a 96.5% GM (98.5% sensitivity and 94.6% specificity) and a 74.4% F1-score (98.5% recall and 60.0% precision) for cell type-/tissue-specific enhancers, and it achieved 95.1% accuracy, a 97.0% GM (99.4% sensitivity and 94.6% specificity) and an 81.5% F1-score (99.4% recall and 69.1% precision) for common enhancers. Additionally, for both cell type-/tissue-specific and common enhancers, PEDLA achieved consistent performance within 20 independent test cell types/tissues sets. These results demonstrate that PEDLA has the properties of invariability (cell type-/tissue-common) and variability (cell type-/tissue-specificity) in enhancer prediction, further suggesting its superior performance consistency across various cell types/tissues.

**Figure 4.**
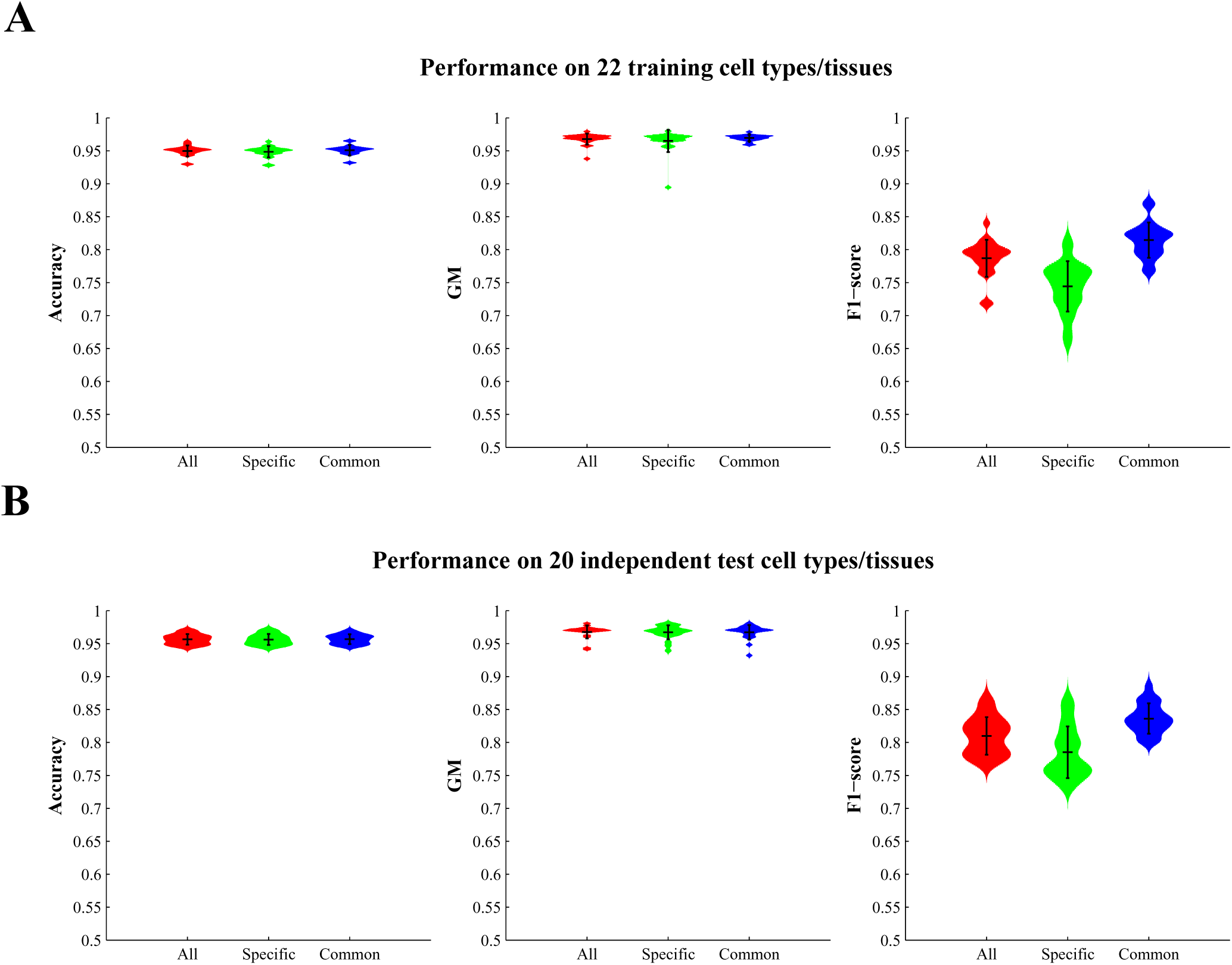
Performance assessment of PEDLA with the best-trained model in multiple human cells/tissues. (A-B) Performance assessment of PEDLA with the best-trained model in the training cell set and test cell set. Three performance indicators, accuracy, GM, and F1-score, were assessed for the PEDLA with the best trained model in the 22 training cell types/tissues (A) and 20 test cell types/tissues (B), independently. The best-trained model was the only one selected in terms of performance from the 200 optimal models 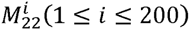 that finished training on 22 training cell types/tissues. All enhancers were classified as “specific” or “common” based on the number of cell types/tissues in which the enhancers occurred. An enhancer that occurred in not more than 4 cell types/tissues was termed specific; otherwise, it was considered common.

### Comparison with the DEEP-ENCODE model

To further illustrate the performance superiority and generalization capabilities of our PEDLA in multiple human cell types, we re-trained and re-tested our PEDLA using the training and testing sets of DEEP-ENCODE model ^44^, respectively. DEEP-ENCODE model, one of the three independent models of DEEP, specialized to identify enhancers using data from ENCODE repository ^44^. For the training sets, DEEP-ENCODE model used GM12878, HepG2, H1-hesc and HUVEC cell lines data coming from ENCODE repository. For testing sets, DEEP-ENCODE model used data from Hela-S3 and K562 cell lines. The feature vector of DEEP-ENCODE model contained 11 histone modification marks, including H2AF.Z, H3K27ac, H3K27me3, H3K36me3, H3K4me1 H3K4me2, H3K4me3, H3K79me2, H3K9ac, H3K9me3, and H4K20me1. To keep consistent with DEEP-ENCODE model, our PEDLA also was trained using these 11 histone modification marks.

For the Hela-S3 cell line, PEDLA predicted 29,308 enhancers, whereas DEEP-ENCODE predictions covered 37,128. For K562 cell line, PEDLA made 32,503 enhancer predictions, whereas DEEP-ENCODE predicted 94,186. The performance comparison analysis was summarized in Tables 6. We found that PEDLA always performed better than DEEP-ENCODE based on the six performance indicators accuracy, sensitivity, GM, F1-score, and validation rate (DHS and P300) in both Hela-S3 and K562 cell lines. Based on specificity and misclassification rate, our PEDLA and DEEP-ENCODE achieved almost consistent results in both evaluated cell lines. Furthermore, we carefully selected thresholds of DEEP-ENCODE to yield comparable numbers of predictions with PEDLA, so as to exclude the bias of different enhancer predictions on performance comparison. We obtained consistent result with aforementioned finding (Table 6). Together, our results illustrated the performance superiority and generalization capabilities of our PEDLA in another completely different training and testing sets, suggesting that our PEDLA consistently achieved much better performance comparing with DEEP-ENCODE model.

**Table 6.**
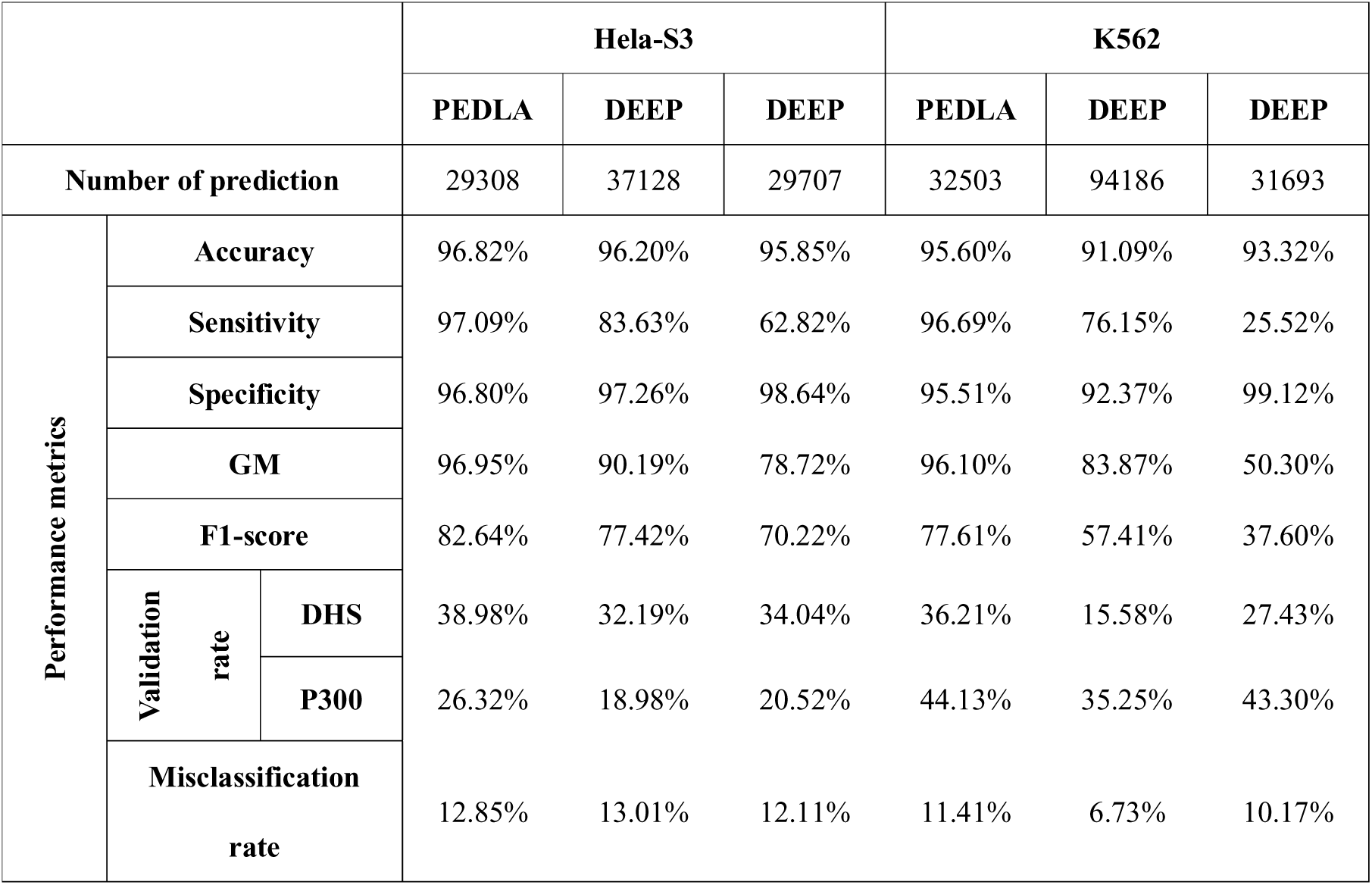
Performance comparisons between PEDLA and DEEP in Hela-S3 and K562 cell lines

## Discussion

In this study, we developed a deep learning-based algorithmic framework named PEDLA (https://github.com/wenjiegroup/PEDLA) to systematically and precisely predict enhancers on a genome-wide scale using heterogeneous types of data. PEDLA has three outstanding characteristics that make it ideal for achieving state-of-the-art performance relative to five existing methods for enhancer predictions. First, PEDLA is capable of learning an enhancer predictor based on massively heterogeneous data to fully capture the universal patterns of enhancers, which makes its enhancer predictions more comprehensive and accurate. In H1 cells, we trained PEDLA using 1,114-dimensional heterogeneous features and 811,036 training samples and demonstrated that PEDLA has three key advantages: limitless training samples, integration of massively heterogeneous data, and an embedded ability to handle highly class-imbalanced data in an unbiased way (Fig. 1). All these merits make PEDLA a general and robust deep learning-based framework for enhancer predictions.

Second, PEDLA can generalize enhancer predictions in ways that are mostly consistent across various cell types/tissues. We extended PEDLA to iteratively learn from multiple cell types/tissues, which is motivated by the core idea of deep learning that uses unsupervised pre-training as initialization of the subsequent supervised fine-tuning. The whole training procedure of PEDLA in multiple cell types/tissues consists of initial training and iterative training (Fig. 2). The initial training procedure adopts a traditional deep learning strategy, whereas the iterative training procedure uses the trained optimal model parameters of a previous cell type/tissue as initialization for the supervised fine-tuning of a subsequent cell type/tissue iteratively. We applied PEDLA to iteratively train on 22 training cell types/tissues, and we found that the learned optimal model manifested superior predicting performance and significant performance consistency in 22 training cell types/tissues and another 20 independent cell types/tissues (Figs. 3–4).

Third, PEDLA is capable of extending to input data of any type and to predictions of any type of functional element/domain. Our PEDLA used nine categories of heterogeneous data in H1 cells as discriminating features to identify enhancers. In fact, any type of data that consists of real values can be served as input features to PEDLA. Additionally, the numbers of input features can range from a few to thousands. Furthermore, PEDLA can be trained on training sets of any number of cell types/tissues but will not result in bloated structures. Most importantly, provided with prepared suitable feature data and constructed accurate positive and negative sets for the task, PEDLA can be extended with almost no modifications to identify any types of functional elements/domains, including promoters, insulators, and repressors.

In our recent study, we presented a novel method, DNN-HMM, for the *de novo* identification of replication domains ^54^ We adopted DNN-HMM model as one unit of our PEDLA. Comparing with DNN-HMM that identified replication timing domains using only Repli-seq data, PEDLA identified enhancers using nine categories of heterogeneous data. Additionally, we illustrated the excellent capability of PEDLA in handling class-imbalanced data unbiasedly, which is a key challenge in enhancer prediction. Furthermore, PEDLA was a flexible framework which could be trained on any number of cell types/tissues and achieved superiorly consistent performance across various cell types/tissues in enhancer prediction. However, DNN-HMM was trained in one cell type, and used the single-cell-trained model to predict replication timing domains in other cell types.

Although PEDLA achieved state-of-the-art performance in enhancer prediction, there is still room for further improvement. First, we carefully selected 1,114-dimensional features derived from nine categories of heterogeneous data in H1 cells; however, it remains challenging to select the comprehensive and discriminating features to identify enhancers; thus, further optimization of the feature set must be continued. Additionally, all these features used by PDELA are two-dimensional (2D). It is expected that integrating three-dimensional (3D) structure information derived from 5C, Hi-C, ChIA-PET and Capture-C will greatly improve the prediction of enhancers, given that 3D chromatin structure is increasingly considered an important regulator of gene expression and that enhancers regulate target genes through promoter-enhancer looping ^55, 56^. A recent study combined Hi-C data with phylogenetic correlations to predict the target genes of distal regulatory elements, such as enhancers, repressors, and insulators ^57^ Another group demonstrated that modelling Hi-C data with their computational method, called graph-based regularization (GBR), greatly improved the prediction of replication and topological domains ^58^. These studies inspired us to further improve PEDLA through the incorporation of 3D structure information. Second, due to the large number of input features and massive training samples, the PEDLA architecture has thousands of weights and millions of connections between units. Whereas training such large networks is time-consuming, progress in algorithm parallelization, distributed computing and GPU acceleration have reduced the training times to a few hours. Third, like many other leading approaches in machine learning, PEDLA based on deep learning is the most data-hungry, requiring thousands and even millions of labelled data to achieve state-of-the-art performance. Thus, reducing the reliance on large amounts of labelled data will greatly improve the applicability of PEDLA. Recently, Brenden M. Lake, *et al*. ^59^ presented a Bayesian program learning (BPL) framework that can learn a large class of visual concepts from only a single example and manifest creative generalization abilities that are mostly indistinguishable from human behaviour. A novel hybrid system combining deep learning and BPL may therefore provide an efficient strategy to dramatically reduce the reliance on large amounts of labelled data. Finally, PEDLA can be directly extended to predict any known functional elements/domains in the genome but it cannot be used to discover completely novel regulatory elements because PEDLA is supervised in nature. Human learning is largely unsupervised because we discover the world by observing it, not by being told what it is in advance ^60^. To achieve human-level learning, it becomes far more important to explore the potential computational model based on deep learning for unsupervised learning in the longer term. Taken together, our work illustrates the power of harnessing state-of-the-art deep learning techniques to consistently identify regulatory elements from massively heterogeneous data across diverse cell types/tissues.

## Materials and Methods

### Feature data

For the construction of PEDLA to predict enhancers in H1 cells, we used nine sets of features, including histone modifications, TFs and cofactors, chromatin accessibility, transcription, DNA methylation, CpG islands, evolutionary conservation, sequence signatures, and occupancy of TF binding sites (TFBS). Features derived from ChIP-Seq data included 27 histone modification marks and 15 TFs and cofactors analysed in the ENCODE Project ^47^ and the Roadmap Epigenomics Project ^48^. Attributes coming from chromatin accessibility and transcription were obtained from DNase-Seq and mRNA-Seq in the Roadmap Epigenomics Project ^48^. Characteristics derived from DNA methylation were obtained from RRBS (Reduced Representation Bisulfite-Seq) in the Roadmap Epigenomics Project ^48^. Features obtained from CpG islands and evolutionary conservation were taken from the CpG island track and the vertebrate phastCons46way track ^61^ in the UCSC Genome Browser ^62^, respectively. A total of 340 sequence characteristics were derived by estimating the frequency of all possible *k*-mers (*k* = 1, 2, 3, 4) in the sequence, and 726 attributes derived from occupancy of TFBS motifs were obtained by scanning TF motif occurrences within open chromatin in the human genome using FIMO ^63^ with a *p*-value < 10^−5^. We extracted 726 TF motifs from Transfac ^64^, Jaspar ^65^ and UniProbe ^66^ and defined open chromatin in the human genome as the merged TFBS-clustered regions across hundreds of cell types ^47^, which covers nearly all accessible regions in the human genome. In total, 1,114-dimensional features were used to predict enhancers with PEDLA in H1 cells (Table S1).

### Data normalization for diverse data types

All 1,114 features were processed and normalized at a 200-bp resolution using a unified RPKM-like measure (reads per kilobase per million) ^67^. The ChIP-Seq mapping reads for the histone modifications, TFs and cofactors and the corresponding inputs were binned into 200-bp intervals, and the RPKM value of each bin was calculated. The DNase-Seq and RNA-Seq mapping reads were also binned into 200 bp intervals, and the RPKM value of each bin was calculated. For two or more replicates of these sequencing data, the RPKM value for each bin was averaged to obtain a single RPKM value. The signals of RRBS, the CpG island, evolutionary conservation, *k*-mers (*k* = 1, 2, 3, 4) sequence characteristics, and occupancies of TFBS motifs in each bin were normalized against the total signal of the whole genome and the length of the bin, similar to the RPKM calculation. Finally, the 1,114 feature signal values were scaled to the range of [0,1].

### Construction of the positive and negative sets

Considering that there are no ‘gold standard enhancers’ that have been experimentally verified across various cell types/tissues, we constructed the enhancer set (positive set) using histone marker H3K27ac, which is viewed as an active enhancer hallmark ^9, 68–70^ and is ubiquitously available across dozens of cell types/tissues. To construct the positive set, we started from the candidate H3K27ac peaks called by MACS ^52^ with a *p*-value < 10^−9^ for each replicate independently. Then, we selected the candidate H3K27ac peaks that were consistently identified in multiple replications. Next, we strictly used the highly reliable H3K27ac peaks, which met the following criteria as described previously ^10^: (1) H3K27ac peaks that were located within 1 kb of all annotated transcription start sites (TSSs) of protein-coding genes annotated by GENCODE (V15) ^50^ were removed. (2) H3K27ac peaks that had a 5’-sequenced expressed sequence tag (EST) with a 5’ end within 2 kb of the H3K27ac peak and spanning an annotated TSS were removed on the basis of annotations of ESTs from the UCSC Genome Browser spliced EST track. (3) H3K27ac peaks that had significant enrichment of both H3K4me1 and H3K4me3 in a 2-kb window centred on the peak were removed. (4) H3K27ac peaks with significant enrichment of H3K4me3 within a 2 kb window centred on the peak were also removed. (5) An H3K4me1 peak had to be present within 2 kb of the H3K27ac peak. (6) The H3K27ac peak had to be at least 2 kb away from all rRNA genes.

Non-enhancers (the negative data set) contained an equal number of promoters and *x* times the number of random genomic loci not annotated as promoters or enhancers. Promoters are defined as 2-kb regions centred on TSSs of protein-coding genes, and random genomic loci were generated with an equivalent length distribution as enhancer regions. For each learning set, we set *x* = 9 to maintain a ratio of 1:10 between positive and negative samples, which is consistent with previous studies ^39, 40, 42, 44^. In the analysis of class imbalance problems, we changed *x* from 1 to 9.

For enhancer and non-enhancer sets of each cell type/tissue, all the 200-bp intervals that overlapped with any enhancers or non-enhancers were compiled to be the final training samples for the development and assessment of our PEDLA. The average length of all enhancer and non-enhancer regions was approximately 2 kb. Therefore, we obtained approximately (1+1+*x*)×*n*×2,000/200 = 10*n*×(2+*x*) training samples for *n* experimentally determined enhancers in the positive set. For H1 cells, we had 5,870 enhancers, and the number of training samples in the H1 cell line was 811,036.

### Training the PEDLA for enhancer predictions

Enhancer prediction with PEDLA, which follows our recently reported method ^54^, involved two stages, namely training and prediction procedures (Fig. S1).

1. Training procedure

1) Read the normalized features and annotated labels as input data.
2) Estimate the initial state probability vector *π* and state transition probability matrix *A* of HMM.
3) Pre-train the DNN layer-by-layer in an unsupervised fashion, then use the pre-training as initialization to fine-tune the weight matrix *W* of the DNN using mini-batch gradient descent in a supervised manner.
4) Estimate the prior probability vector *P* = {*p*(*q*_*t*_ = *S*_*j*_)} = {*n*(*S*_*j*_)/*n*}, where *n*(*S*_*j*_) is the number of samples labelled with state *S*_*j*_, and *n* is the total number of all samples.
2. Prediction procedure

1) Calculate the output of the DNN to estimate the posterior probability of each state based on the weight matrix *W*.
2) Divide the posterior probability by the prior probability *P* to obtain the observation probability distribution *B* of HMM.
3) Use the Viterbi algorithm to predict the final state of unlabelled data based on the triple (*π*, *A*, *B*).

### Performance assessment of PEDLA

Our PEDLA employed a threshold-free decision mechanism and could automatically make the optimal decision to output the most reasonable enhancer predictions. The threshold-dependent receiver operation curves (ROCs) and corresponding the area under the curve (AUC) cannot used to assess the performance of PEDLA. To compare the performance across these methods, we computed the following three performance indicators used in our recent study ^54^.

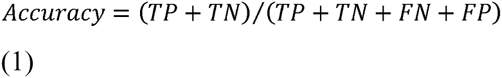

where TP indicates true positives, FP indicates false positives, TN indicates true negatives, FN indicates false negatives, and

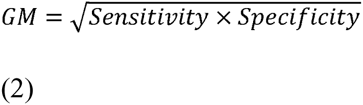

where GM is the geometric mean of sensitivity and specificity, Sensitivity = TP/(TP+FN) and Specificity = TN/(TN+FP), and

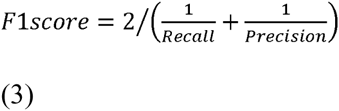

where the F1 score is the harmonic mean of recall and precision, Recall = Sensitivity and Precision = TP/(TP+FP).

For PEDLA and other supervised methods, we used 5-fold cross-validation to obtain unbiased estimates of the three performance indicators. For the unsupervised methods, ChromHMM and Segway, we obtained these three performance indicators by comparing their enhancer predictions with the labels of the positive and negative sets.

### Selecting optimal structure of PEDLA

To determine the optimal structure of hidden layers for PEDLA, we tuned the structure of hidden layers by fixing the number of hidden layers to 1 or 2 and the number of hidden units in each hidden layer to 10, 50, 100 or 500. We made enhancer predictions using PEDLA for diverse structures that are composites of various hidden layers and hidden units of each layer. We computed the three performance indicators using 5-fold cross-validation and compared them between the structure models of PEDLA to select the optimal structure model of PEDLA in terms of accuracy (Table S3).

### Validation of enhancer predictions

To validate the enhancers predicted by PEDLA and other existing methods, we calculated the percentage of the predicted enhancers that overlapped with experimental data on enhancer markers (the validation rate) or TSSs of protein-coding genes (the misclassification rate). The enhancer markers included distal DNase-I hypersensitivity sites (DHSs), binding sites of p300 and TFs such as NANOG, OCT4 and SOX2. For H1 cell lines, the DHSs were derived from an ENCODE study (http://ftp.ebi.ac.uk/pub/databases/ensembl/encode/integration_data_jan2011/byDataType/openchrom/jan2011/fdrPeaks/) ^47^, and p300, NANOG, OCT4 and SOX2 binding sites were obtained from a recent study ^40^. For Hela-S3 and K562 cell lines, DHSs and p300 were obtained from the study of DEEP ^44^ TSS annotations were extracted from the GENCODE annotations (V15) ^50^. Predicted enhancers overlapping with a window of −100 to +100 bp centred at a distal enhancer marker that was greater than 5 kb away from the nearest TSS were classified as ‘‘validated’’, whereas predicted enhancers located within 2.5 kb of a TSS were classified as “misclassified”.

## Availability

The source code of our PEDLA method can be freely accessed at https://github.com/wenjiegroup/PEDLA.

## Acknowledgements

We wish to thank the ENCODE Project and Roadmap Epigenomics Project for making their data publicly available. This work was supported by grants from the Major Research plan of the National Natural Science Foundation of China (No. U1435222), the Program of International S&T Cooperation (No. 2014DFB30020) and the National High Technology Research and Development Program of China (No. 2015AA020108).

## Author Contributions

W.S. conceived the study. W.S. and X.B. designed all experiments. WS drafted the manuscript. FL wrote the programs, analyzed the results. HL and CR and helped in analysis and discussion, gave useful comments. All authors read and approved the final manuscript.

## Supplementary Information

Supplementary information accompanies this paper at http://www.nature.com/scientificreports.

Competing financial interests: The authors declare no competing financial interests.

